# Object-based attention prioritizes working memory contents at a theta rhythm

**DOI:** 10.1101/369652

**Authors:** Benjamin Peters, Jochen Kaiser, Benjamin Rahm, Christoph Bledowski

## Abstract

Attention selects relevant information regardless of whether it is physically present or internally stored in working memory. Perceptual research has shown that attentional selection of external information is better conceived as rhythmic prioritization than as stable allocation. Here we tested this principle using information processing of internal representations held in working memory. Participants memorized four spatial positions that formed the endpoints of two objects. One of the positions was cued for a delayed match-non-match test. When uncued positions were probed, participants responded faster to uncued positions located on the same object as the cued position than to those located on the other object, revealing object-based attention in working memory. Manipulating the interval between cue and probe at a high temporal resolution revealed that reaction times oscillated at a theta rhythm of 6 Hz. Moreover, oscillations showed an anti-phase relationship between memorized but uncued positions on the same versus other object as the cued position, suggesting that attentional prioritization fluctuated rhythmically in an object-based manner. Our results demonstrate the highly rhythmic nature of attentional selection in working memory. Moreover, the striking similarity between rhythmic attentional selection of mental representations and perceptual information suggests that attentional oscillations are a general mechanism of information processing in human cognition. These findings have important implications for current, attention-based models of working memory.

---

Recent work has shown that allocation of attention toward information in the environment is not continuous, but rhythmical and discrete in time (VanRullen, 2018). For example, when participants are cued to attend a spatial location, target detection rate varies over time since cue onset in a rhythmic fashion at frequencies in the theta and alpha ranges (approx. 4-12 Hz). Such behavioral oscillations have been observed in a variety of perceptual tasks involving visual or auditory target detection (Dugué, McLelland, Lajous, & VanRullen, 2015; Ho, Leung, Burr, Alais, & Morrone, 2017; Landau & Fries, 2012; VanRullen, Carlson, & Cavanagh, 2007) or priming (Huang, Chen, & Luo, 2015). Moving beyond spatial attention, Fiebelkorn et al. (2013) demonstrated that visual attention selects perceptual objects at a theta rhythm. They presented pairs of objects and cued the endpoint of one of them as the likely site of an upcoming target (Egly, Driver, & Rafal, 1994). To measure object-based attention independently of visuospatial attention elicited by the cue, they compared performance at the uncued endpoint of the same object as the cued position (i.e., the same-object position) with performance at an endpoint of the other, unattended object (i.e., the different-object position). Perceptual processing was organized rhythmically as performance oscillated at a theta rhythm and in an anti-phase relationship between both positions, suggesting rhythmic object-based prioritization. Together, results from different perceptual studies indicate that rhythmical prioritization constitutes a general attentional mechanism of processing external information.

According to current models, attention is an integral part of working memory (WM), as allocation of attention supports the “working” part of memory by providing the means to keep memories active and amenable to direct access and to flexibly prioritize currently relevant pieces of information among those maintained in WM (Cowan, 1988; Oberauer, 2002). For example, attention can be directed to internally stored positions in WM via a “retro-cue” during the retention interval. Similar to perception, this leads to faster reaction times for memory probes presented at the cued than uncued positions (Griffin & Nobre, 2003; Landman, Spekreijse, & Lamme, 2003). It is therefore possible that rhythmic attention should be detected not only for external stimuli but also for information maintained in WM.

Recently, we have demonstrated that the principles of perceptual object-based selection equally apply to objects in WM (Peters, Kaiser, Rahm, & Bledowski, 2015). Specifically, we found shorter reaction times when participants shifted attention to a memorized position located on the same-object representation as compared to a position located on the different-object representation. This task was particularly well suited to study the effects of attention in WM. Assuming that spatial attention was directed to the currently cued position, differences between equidistant same- and the different-object memory positions could be attributed to the objects encoded in WM and thus unequivocally dissociated from purely visuospatial attention. Thus, this manipulation allowed to dissociate object-based attention in WM (same-versus different-object position), which represented our study question, from spatial attention (cued versus uncued memory positions) where perceptual and WM effects can be intermixed. Here, we employed a version of this paradigm by varying the time interval between attentional cue and probe with a high temporal resolution to test whether attention to internal object representations in WM is also modulated rhythmically. We hypothesized that the latency of object-based selection in WM should oscillate at a theta rhythm with an anti-phase relationship between same- and different-object positions.

## Methods

### Participants

30 participants (16 female; mean age = 22.1 years) with normal or corrected-to-normal vision were recruited from the Goethe-University Frankfurt and the Fresenius University of Applied Sciences Frankfurt and gave written informed consent. We excluded two participants with correct-response rates below 60%. Participants were either remunerated or received course credit. The study was approved by the local ethics committee.

### Task and Stimuli

Participants viewed four gray dots (2500 ms; size in visual angle: 0.37°) that were grouped by “banana-shaped” light gray regions to form the endpoints of two objects (Figure 1). Spatial positions and objects were presented on an invisible circle with a radius of 8° visual angle around fixation. Neighboring memory positions were separated by 60°, 90°, or 120° angular distance on the invisible circle.

**Figure 1.**
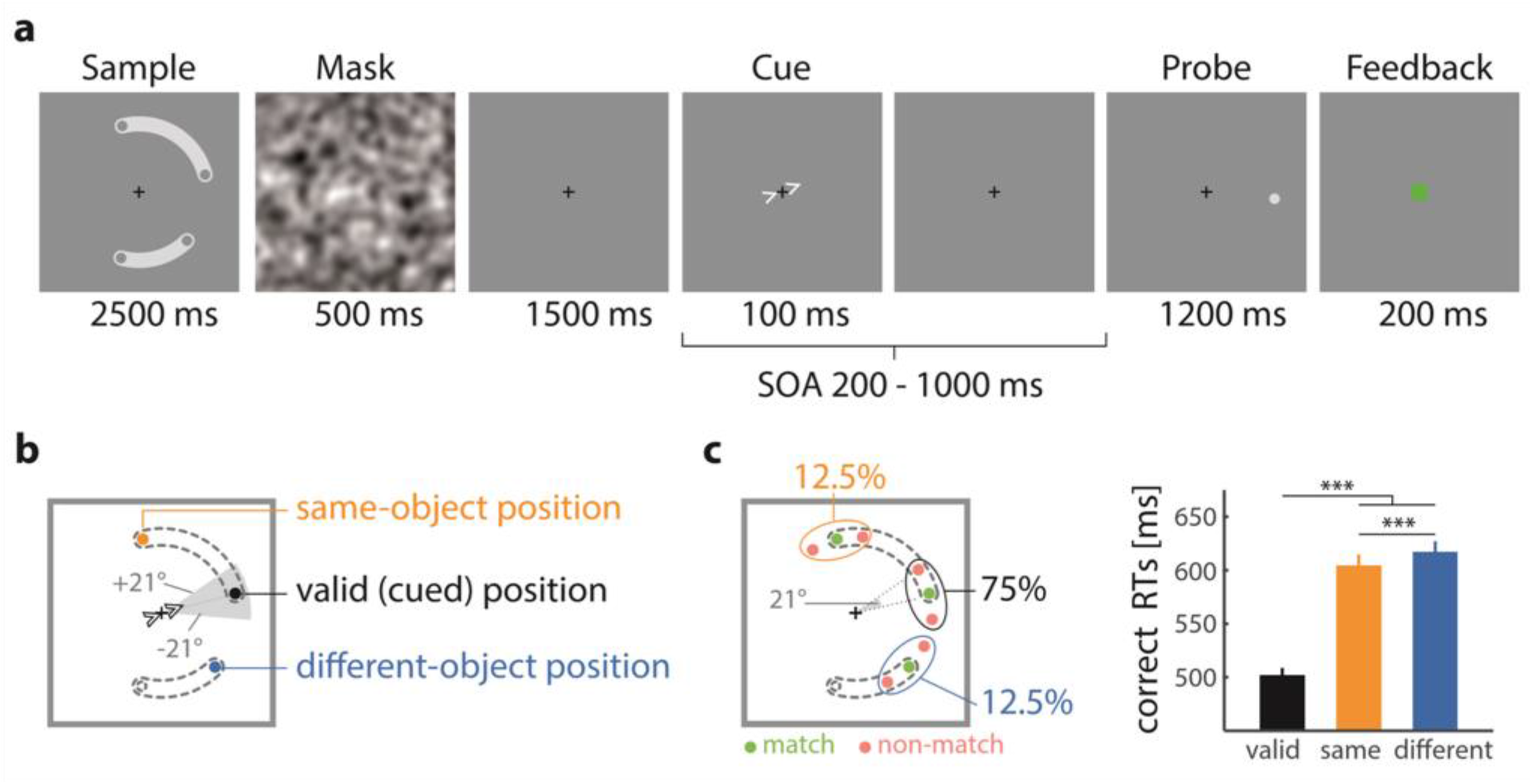
Experimental paradigm. **a** Participants memorized the exact locations of the endpoints of the two objects. Object-length and position varied randomly across trials. During retention, a spatial cue indicated the endpoint location which would most likely be probed. **b** Memory positions. To ensure that participants selected information from working memory upon cue presentation and to prevent the cue from revealing the memorized position, the cue pointed to the vicinity (±21°) of the cued memory position but was uninformative about the match/non-match decision at the cued position. **c** The cued position (black) was probed in 75%, the same-object position (yellow) in 12.5% and different-object position (blue) in 12.5% of all trials. Probes were either matching the corresponding memory position (green) or displaced by ±21° (red). **d** Reaction times (RTs) of correct responses for the three probe conditions. Lines indicate standard error of the mean. SOA stimulus-onset-asynchrony. *** *p* <. 001, paired *t*-test.

The location of the gray dots and the groupings varied randomly across trials. After encoding, a visual mask was presented (500 ms). During the retention interval a central cue (100 ms) pointed to a position in the vicinity of one of the four memory positions to avoid revealing the exact memory position. The cue’s direction was drawn from a uniform distribution (−21° to +21°, centered on the cued memory position). Stimulus onset asynchronies (SOAs) between cue and probe were equidistant from 300 ms to 1000 ms in steps of 33.3 ms (two frames on a 60-Hz monitor). In addition, two shorter SOAs (200 ms and 267 ms) were used at time points before attentional selection in WM has been shown (Souza & Oberauer, 2016). Matching probes were presented at the exact spatial location as during encoding. Non-matching probes were displaced from the probed memory position either clockwise or counterclockwise by 21°. The probe could appear either at the cued memory position, at the uncued memory position on the same object or at the spatially nearest memory position on the uncued object. Participants gave their response via left (match) or right (non-match) mouse button click. Probes were presented for 1200 ms. Performance feedback was presented (200 ms) consisting of a green (correct) or red central square (incorrect or reaction time (RT) > 1200 ms).

### Procedure

Each participant conducted 60 trials of the valid cue condition, 10 trials of the same-object condition, and 10 trials of the different-object condition for each of the 23 SOAs. The resulting 1840 trials were separated into 46 blocks of 40 trials each that were performed in four separate sessions within two weeks. Sessions lasted approximately 1.5 h each and were always conducted on separate days.

### Data analysis

RTs for correct responses were averaged in two adjacent time bins using a sliding time window approach for the analysis of the time courses (see Fiebelkorn et al., 2013) before entering analyses or in figures. To obtain the frequency spectrum of the time course of object-based attention in WM (i.e., Fig. 2c) we analyzed the difference of RTs in the same-object condition and different-object condition for the SOAs with equidistant temporal spacing (excluding the first two SOAs). Each participant's difference time-course was detrended using a second order polynomial and subsequently Fourier transformed. Amplitude values at frequencies from 1.5 Hz to 13.5 Hz (in steps of 1.5 Hz) were averaged across participants to obtain the mean amplitude spectrum. To obtain the phase relationship of the 6-Hz oscillations at the same- and different-object positions (i.e., Fig. 3), we performed the Fourier transform for the same- and different-object conditions separately as described above (see Supplementary Figure 3). We then computed the angular difference between the phase angles of the 6-Hz oscillations of each condition. The angular difference was then projected onto the unit circle in the complex plane and averaged across participants. The length and the angle of the resulting vector corresponded to the phase-locking value (PLV, Lachaux, Rodriguez, Martinerie, & Varela, 1999) and the mean phase difference, respectively. We obtained non-parametric estimates of the probability of the observed data under the null hypothesis.

#### Statistics

For each of 5000 permutation samples, each participant’s individual time-course was shuffled before entering the analysis as described above (i.e., before applying the sliding-time window). This resulted in one mean amplitude spectrum for the object-based attention effect, one PLV, and one mean phase difference between the same- and different-object condition for each permutation sample. Individual frequency *p*-values in the amplitude spectrum were corrected for number of frequency bins to control the false discovery rate at 5% (Benjamini & Hochberg, 1995).

**Figure 2.**
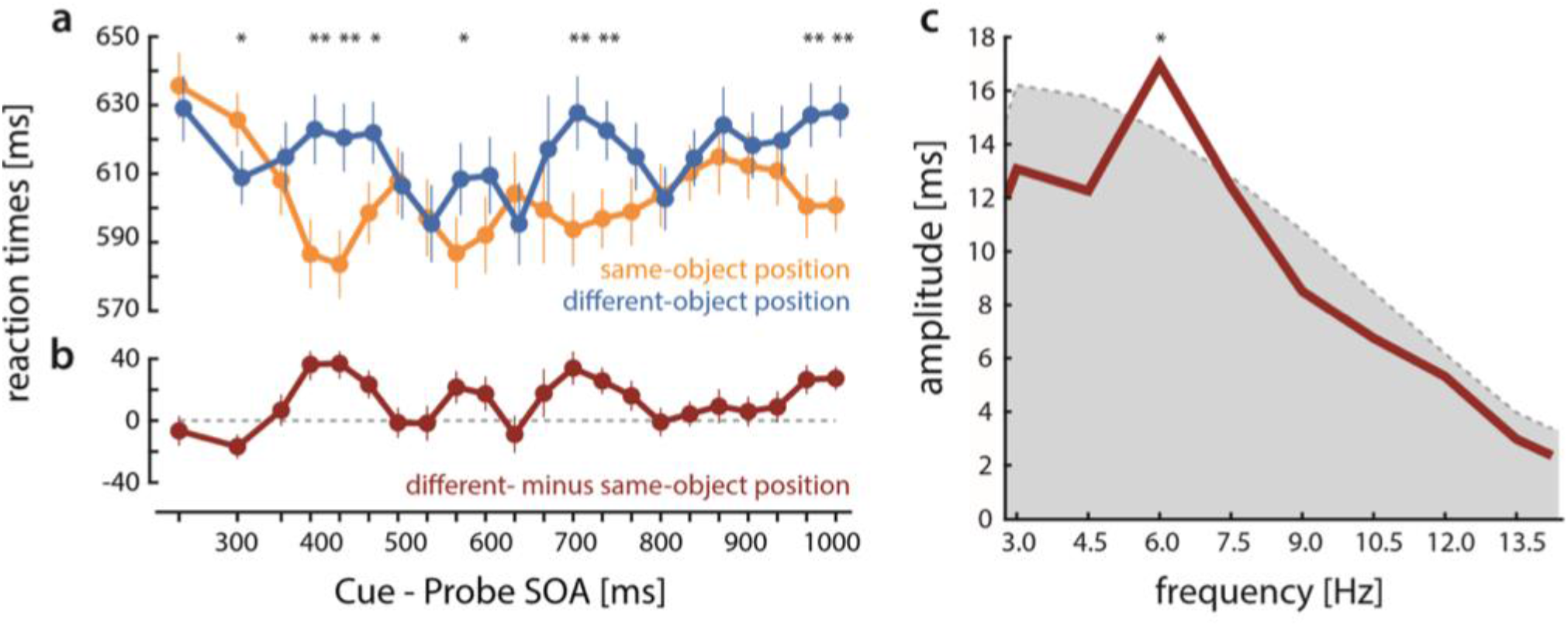
Results. **a** Reaction times (RTs) to the probe as a function of time after the onset of the attentional cue (SOA: stimulus-onset-asynchrony) for the memorized same-object position (yellow) and the memorized different-object position (blue). Lines indicate within subject standard errors (Morey, 2008) * *p* < .05, ** *p* < .01. **b** Difference in RTs between different-object condition and same-object condition (i.e., object-based attention). Vertical lines indicate within-subject standard errors (Morey, 2008). **c** Fourier amplitude spectrum of the time-course of object-based attention in working memory (RTs of different-object condition minus same-object condition), shaded area represents 95^th^ percentile of permutation null-distribution * *p* <. 05, FDR-corrected (Benjamini & Hochberg, 1995).

**Figure 3.**
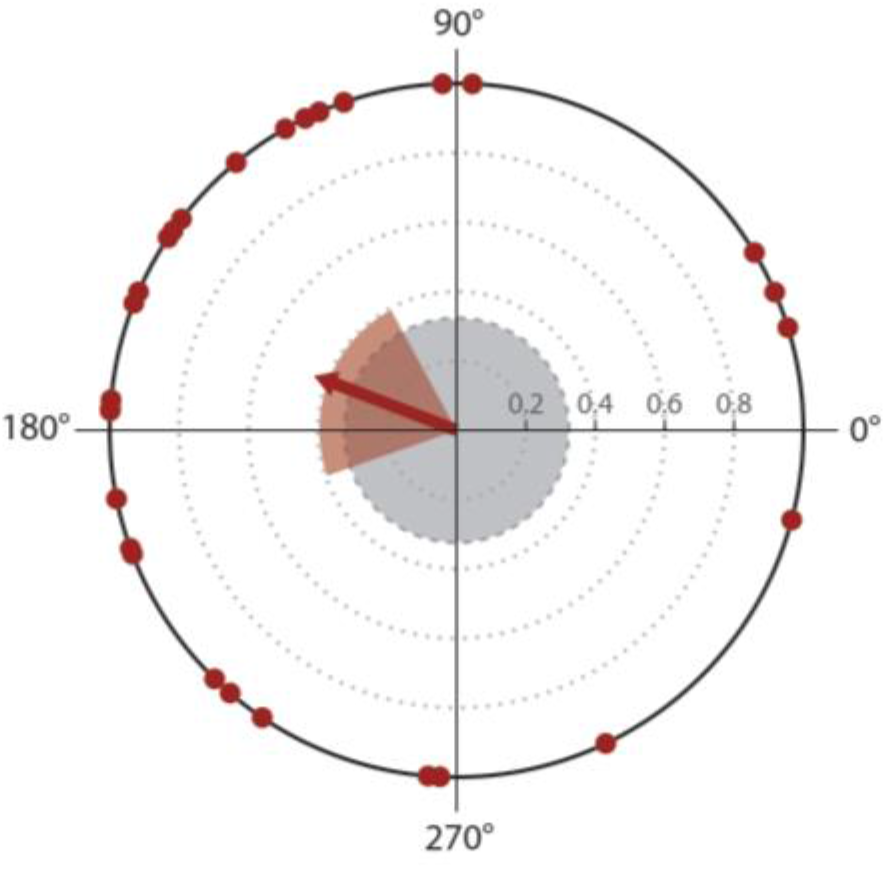
Phase relationship of 6Hz oscillations for the same- and different-object condition in working memory. Dots indicate participants’ individual phase differences plotted on the unit circle. Length of the resultant vector corresponds to the phase-locking value (PLV). Red shaded area indicates 95% confidence interval of the phase difference. Gray shaded area indicates 95^th^ percentile of the permutation null distribution of PLVs.

## Results

We replicated our previous findings of object-based attention in WM (Peters et al., 2015) with faster RTs to a probe presented at memory positions located on the currently attended object compared with equidistant positions on the unattended object (*F*(1,27) = 17.42, *p* < .001, η_p_^2^ = .39, mean difference ∆t = 13 ms, Figure 1d). We also observed that participants showed faster RTs (*t*(27) = 20.96, *p* < .001) and higher accuracy (*t*(27) = 15.27, *p* < .001) for the cued memory position compared to the two uncued memory positions consistent with the allocation of focal attention at the cued position (Egly et al., 1994; Fiebelkorn et al., 2013; Peters et al., 2015).

Assessing the temporal dynamics of object-based attention in WM, we found that the object-based attentional benefit varied as a function of the temporal lag between cue and probe (interaction: time × condition: *F*(21,567) = 2.35, *p* < .001, η_p_^2^ = .08). The benefit emerged at approximately 383 ms after cue onset (*t*(27) = 3.59, uncorrected *p* = .001, ∆t = 36 ms), indicating that, by this time, object-based attention in WM had co-activated the same-object position (Figure 2a). Crucially, we found clear evidence for attentional rhythmicity in WM. In particular, after the initial build-up of the object-based attentional benefit in WM at 383 ms, it vanished at 483 ms (*t*(27) = 0.14, *p* = .89, ∆t = 1 ms) and subsequently re-occurred periodically with peaks at 550 ms, 683 ms, and 950 ms (Figure 2a). This shows that object-based attention in WM oscillated in time even though the probability of a target appearance was identical for both uncued memory positions throughout the cue-probe interval. Fluctuations of object-based RT benefits across time were further supported by a significant interaction between time and condition even when the first three temporal lags, during which object-based attention was in the process of building up, were excluded (*F*(18,486), *p* = .015, η_p_^2^ = .07). When calculating an amplitude spectrum of frequencies from 1.5 Hz to 13.5 Hz (Figure 2b), we observed a significant peak at 6 Hz (*p* < .05, FDR corrected). This reveals that object-based attention in WM fluctuated in the theta range, corroborating the frequencies of rhythmic attention observed in perception.

In contrast to the oscillatory same-object benefit we did not observe any rhythmic fluctuation at the validly cued location (see Supplementary Analysis 1 for a detailed analysis of behavioral responses at the cued memory position).

A rhythmic same-object benefit could have been driven by fluctuations at either the same- or different-object position alone. However, if behavioral oscillations resulted from rhythmic object-based prioritization in WM, the speed of access for the cued and uncued object should be anti-correlated, as previously shown for visual spatial and object-based attention (Fiebelkorn et al., 2013; Landau & Fries, 2012; Song, Meng, Chen, Zhou, & Luo, 2014). We found that the 6 Hz rhythms of the time course of RTs in the same- and different-object conditions were significantly phase-locked (phase-locking value = .40, *p* = .011). Moreover, the mean phase angle difference between the two memory conditions was 159° (95% confidence interval from 119° to 199°, Figure 3) revealing an anti-phase relationship of both oscillations. Notably, reaction times at the different-object position were never faster than at the same-object position, indicating alternating periods of prioritization of the same-object position and periods of equal prioritization of the same- and different-object positions, as previously observed for object-based attention in perception (Fiebelkorn et al., 2013).

The paradigm was designed to study the effect of object-based attention on reaction times. However, we also analyzed accuracies for which we observed similar albeit slightly weaker effects (see Supplementary Analysis 2) and we found no evidence for a speed accuracy trade-off (see Supplementary Analysis 3).

## Discussion

The present study demonstrated a rhythmic modulation of attention to information that is not physically present but retained internally in WM. This suggests that attentional oscillations extend beyond perception and thus represent a general mechanism of cognitive information processing. Specifically, we found that object-based attention in WM oscillated rhythmically at a rate of 6 Hz.

When a part of an object is cued, attention is directed to it, which leads to strong behavioral benefits for that object part. In addition, object-based attention involves a spread of attention to all uncued parts of that object, which can be measured by a slight behavioral advantage for these uncued parts of the same object compared to equidistant parts of other objects (Egly et al., 1994). This object-based attention effect has previously been demonstrated in WM (Peters et al., 2015). Here, we found that object-based attention in WM oscillated at a theta rhythm. In particular, we observed a preferential processing that extended from the cued location to the same-object location leading to faster reaction times to probes at the same-object position. This object-based benefit, however, alternated periodically with time windows when probes at the different-object location were responded to with the same speed as those at the same-object location. These results suggest that attention never literally switched to the different-object location (which would entail phases of shorter RTs to different-object positions), but remained on the cued object. In full accordance with this view, we did not observe any fluctuations of RT at the cued position which may indicate that attention persisted there throughout the interstimulus interval (see Supplementary Analysis 1 for performance at the cued position). These findings would not be expected if at each time-point either the same- or the different-object location was attentionally focused, but never both. Together, this suggests that attentional prioritization alternates rhythmically between two phases: in the first, object-based phase, prioritization spreads from the cued position across the whole cued object to the same-object position while concurrently leading to a deprioritization of the other, different-object position (Egly et al., 1994; Fiebelkorn et al., 2013, Peters et al., 2015). In the second phase, attentional prioritization still remains on the cued memory position but does not expand across the complete cued object. Rather, it is symmetrically distributed between the same- and different-object position (Fiebelkorn et al., 2013), leading to similar RTs there.

In perception, evidence for waxing and waning of attention has been referred to as ‘rhythmic attentional sampling’ (VanRullen, 2018; Fiebelkorn & Kastner, 2019). In particular, it has been suggested that objects and locations are sampled in alternation by attention (Landau & Fries, 2012; Fiebelkorn et al., 2013). Regarding object-based attention Fiebelkorn et al. (2013) found that it prioritizes objects at a theta rhythm. Similar to the present study, Fiebelkorn et al. (2013) found rhythmic reweighting of attention to objects, whereby attention never fully switched to the other object. This was evidenced by behavioral performance that was never better at the different-object position than at the same-object position. We observed a strikingly similar pattern of theta-rhythmic object-based attention at the same- and different-object locations that were not perceptually present but held in WM. These findings underscore the notion of shared attentional mechanisms between perception and WM (Gazzaley & Nobre, 2012). In particular, rhythmic attentional prioritization ^1^ therefore constitutes a more general mechanism of information selection and processing than hitherto assumed.

What is the functional role of rhythmic attention? In perception, it has been suggested that periodical reweighting of attention (Buschman & Kastner, 2015) may support environmental scanning and thus constitutes a mechanism of exploration behavior (Fiebelkorn & Kastner, 2019) that facilitates selection of new, so far unselected information in the environment despite attention being focused at the same time. A scanning mechanism would in particular reweight priorities of currently unselected but potentially relevant objects or locations (Jensen, Bonnefond, & VanRullen, 2012). In accordance with this view, we observed oscillations of performance in response only to unselected (i.e., uncued) memory positionsOnly these positions required shifting of the focus of attention to the probed positions for the subsequent response. In contrast, the cued memory position was selected directly after cue presentation and remained in the focus of attention until probe presentation. This suggests that the theta-rhythmic prioritization modulated processes that are involved in selecting the probed memory position and shifting the focus of attention (e.g., Souza, Rerko, & Oberauer, 2016).

Research has shown that attention to WM contents is fully deployed approximately 300 to 500 ms after the onset of an attentional retro-cue, similar to the finding in our study (Gressmann & Janczyk, 2016; Schneider, Mertes, & Wascher, 2016; Souza & Oberauer, 2016; Tanoue & Berryhill, 2012; van Moorselaar, Gunseli, Theeuwes, & Olivers, 2015). However, until now, it has been assumed that attention in WM is allocated stably after its initial deployment (Souza & Oberauer, 2016). Testing with higher temporal resolution than previous studies and using a paradigm that allowed us to dissociate object-based attention in WM from externally or internally directed visuospatial attention, we have demonstrated for the first time that object-based attention in WM is not stably allocated but oscillates rhythmically.

What are the neuronal sources subserving theta-rhythmic prioritization in WM? Attentional networks including fronto-parietal (Busch & VanRullen, 2010; Fiebelkorn, Pinsk, & Kastner, 2018) and posterior sites in the primate visual cortex (Landau, Schreyer, Van Pelt, & Fries, 2015; Spyropoulos, Bosman, & Fries, 2017) have been identified to convey neural theta oscillations. These theta rhythms are modulated by attention (Spyropoulos et al., 2017) and their phase predicts behavioral performance in visual detection tasks (Busch & VanRullen, 2010; Fiebelkorn et al., 2018; Landau et al., 2015). Attention in perception and WM have been shown to rely on similar neuronal substrates (Bledowski, Kaiser, & Rahm, 2010; Gazzaley & Nobre, 2012; Peters et al., 2015). This suggests that similar neural substrates may also be involved in the generation of the theta-rhythmic prioritization that we observed in WM. Specifically, theta-mediated fronto-parietal attention systems (Cavanagh & Frank, 2014) may have prioritized object representations putatively stored in posterior regions. Detectability of behavioral oscillations necessarily implies that both the phase of the underlying theta oscillation and the order in which objects were prioritized was sufficiently consistent across trials. As systematic prioritization of objects was only possible by processing the attentional cue, the existence of theta oscillations further suggests that these were phase-reset or initiated by cue presentation.

To summarize, the present psychophysical study provided evidence for a basic mechanism of human cognition that shapes information processing by prioritizing information at a theta rhythm, regardless of whether this information is physically present or held as internal representations in WM.

## Supplementary material

### Supplementary analyses

Supplementary analysis 1: Responses to the cued (valid) position

Supplementary analysis 2: Object-based attention effects on accuracy

Supplementary analysis 3: Analysis of a possible speed-accuracy tradeoffs

### Supplementary Figures

Supplementary Figure 1: Accuracy for the three probe conditions.

Supplementary Figure 2: Temporal dynamics of object-based attention effects on accuracy

Supplementary Figure 3: Fourier amplitude spectrum of correct RTs for individual conditions.

Supplementary Figure 4. Temporal dynamics of RTs to probes at the cued memory position

Supplementary Figure 5. Temporal dynamics of accuracy to probes at the cued memory position

### Supplementary References

### Supplementary analyses

#### Supplementary analysis 1: Responses to the cued (valid) position

We analyzed the reaction times and accuracy to probes at the cued memory position (Supplementary Figure 4 and Supplementary Figure 5). Consistent with previous findings (Egly et al., 1994; Fiebelkorn et al., 2013), responses were faster (*t*(27) = 20.96, *p* < .001) and more accurate (*t*(27) = 15.27, *p* < .001) for the cued memory position compared to the two uncued (i.e. same and different) memory positions (Figure 1d and Supplementary Figure 1). Reaction times but not accuracy was significantly modulated as a function of the temporal lag between cue and probe (response times: *F*(21, 567) = 23.24, *p* < .001, η_p_^2^ = .46; accuracy: *F*(21,567) = 0.85, *p* = .659, η_p_^2^ = .03). There was no significant peak in either amplitude spectrum, suggesting no rhythmic modulation of performance at the cued location in the range of 1.5 to 13.5Hz, consistent with the notion that attention never fully left the cued object.

It is important to note that the experiment was optimized for the detection of reaction time oscillations at the same- and different-object position. In particular, to require participants to focus attention to the cued memorized position, the experimental design had to be uninformative to the match/non-match decision at the cued location. The cue therefore pointed only near the cued memory position (see Figure 1 b) and this spatial jittering across trials may have obscured any nuanced systematic effects to probes at the cued location.

#### Supplementary analysis 2: Object-based attention effects on accuracy

We repeated all analysis steps using accuracy rates instead of RTs as dependent variable. In accordance with object-based attention, accuracy at the same-object position was significantly higher than at the different-object position (*F*(1,27) = 24.2, *p* < .001, η_p_^2^ = .47, Supplementary Figure 1). However, there was neither a main effect of time (*F*(21,567) = 0.9, p = .62, η_p_^2^ = .03) nor a significant modulation of the object-based accuracy benefit across time (time × condition interaction *F*(21,567) = 0.96, *p* = .52, η_p_^2^ = .03, Supplementary Figure 2). The frequency-resolved amplitude spectrum showed a visibly identifiable peak at 6 Hz, which did not reach statistical significance (uncorrected *p* = .069, all other frequencies *p* > .30). Thus, while rhythmic attentional prioritization affected the latency of accessing memorized positions in WM, we found no clear evidence for a modulation of their fidelity.

#### Supplementary analysis 3: Analysis of possible speed-accuracy tradeoffs

To test whether there was a speed-accuracy trade-off in the data, we correlated the time courses of correct reaction times and accuracy for each participant and checked whether these correlations were statistically different from zero. However, we did not observe significant associations between RTs and accuracy in any of the three conditions (valid condition: mean correlation across participants = −.01, *t*(27) = 0.73, *p* = 0.469; same-object position: −0.03, *t*(27) = −0.53, *p* = 0.604; different-object position = 0.04, *t*(27)=0.734, *p* = 0.469). Hence, we found no evidence for a speed-accuracy tradeoff which could have underlain the oscillatory RT effects.

### Supplementary Figures

**Supplementary Figure 1:**
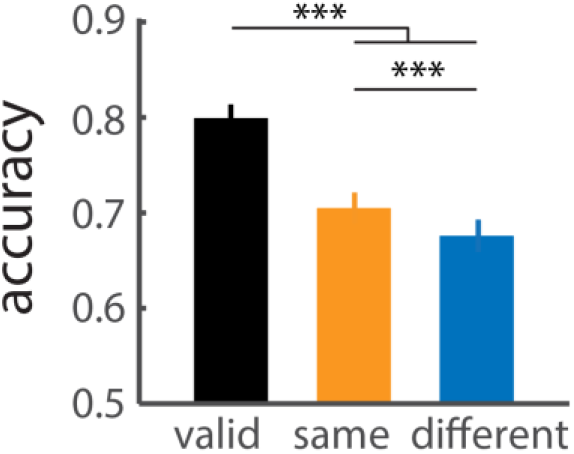
Accuracy for the three probe conditions. Lines indicate standard error of the mean. *** *p* < .001.

**Supplementary Figure 2:**
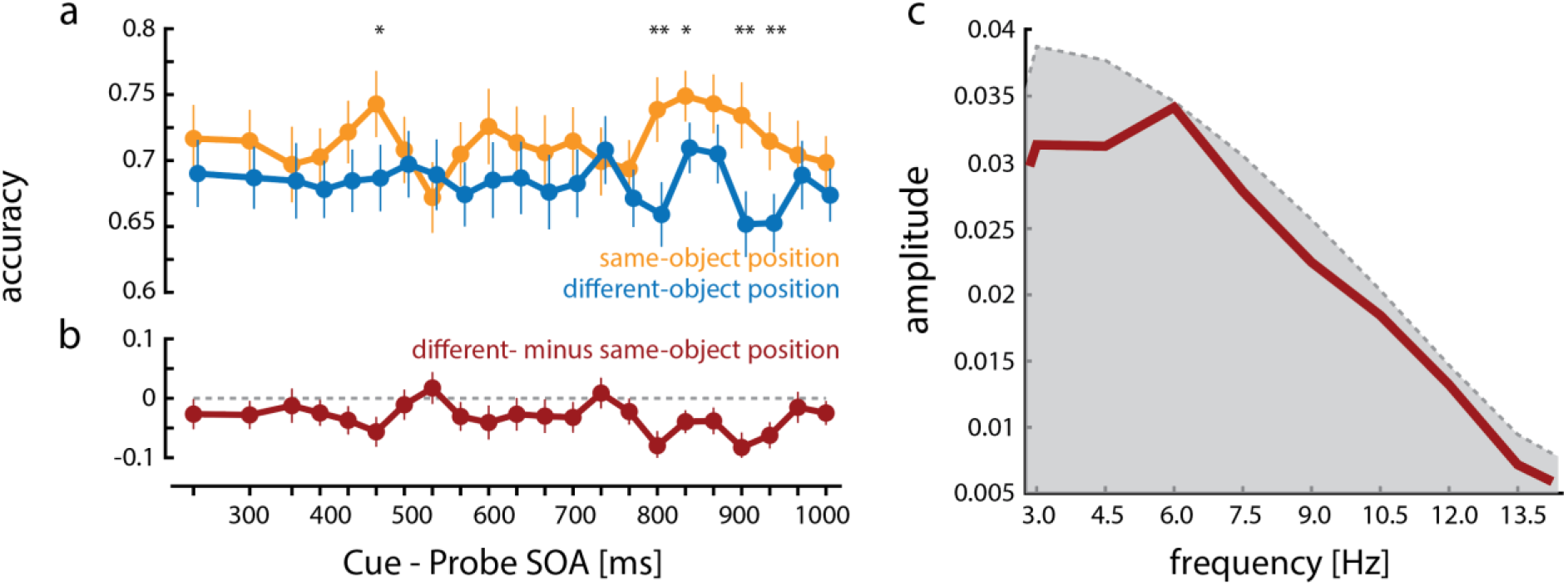
Temporal dynamics of object-based attention effects on accuracy. **a** Accuracy to the probe as a function of time after the onset of the attentional cue (SOA: stimulus-onset-asynchrony) for the memorized same-object position (yellow) and the memorized different-object position (blue). Lines indicate within-subject standard errors (Morey, 2008) * *p* < .05, ** *p* < .01. **b** Difference in accuracy between different-object condition and same-object condition. Vertical lines indicate within-subject standard errors (Morey, 2008). **c** Fourier amplitude spectrum of the time-course of object-based attention in working memory (accuracy of different-object condition minus same-object condition), shaded area represents the 95^th^ percentile of the permutation null-distribution.

**Supplementary Figure 3:**
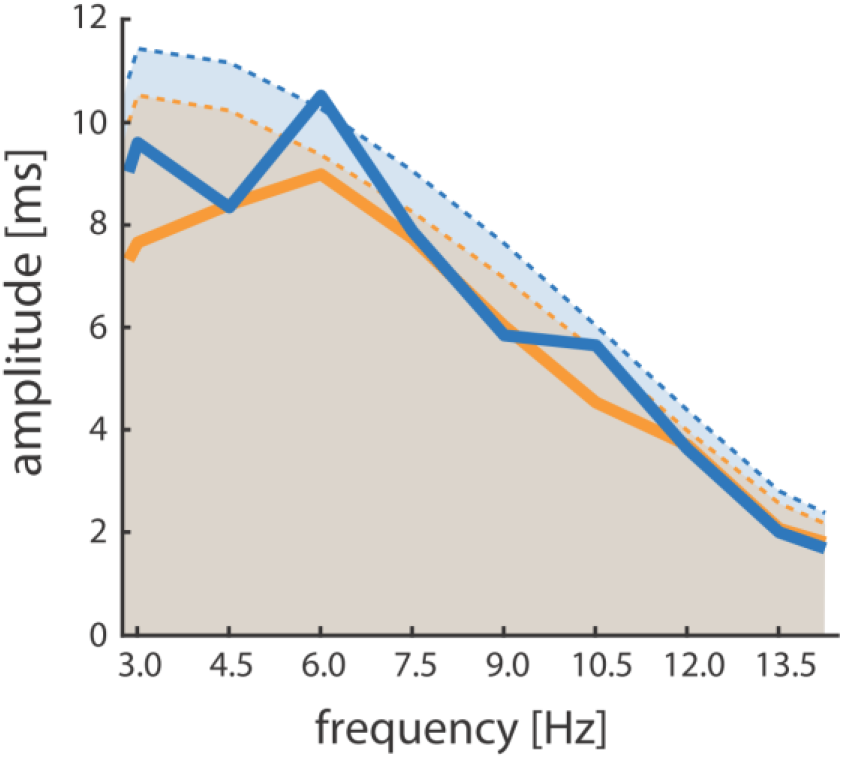
Fourier amplitude spectrum of correct RTs for individual conditions. Same-object position (yellow) and different-object position (blue). Uncorrected p-values for peaks at 6Hz: *p* = .028 (different-object condition) and p = .129 (same-object position). Note: Individual conditions entail only half the data each compared to the object-based attention effect and therefore entail lower signal-to-noise ratios (Figure 2c). Shaded areas represent the 95th percentile of the permutation null-distribution per condition.

**Supplementary Figure 4.**
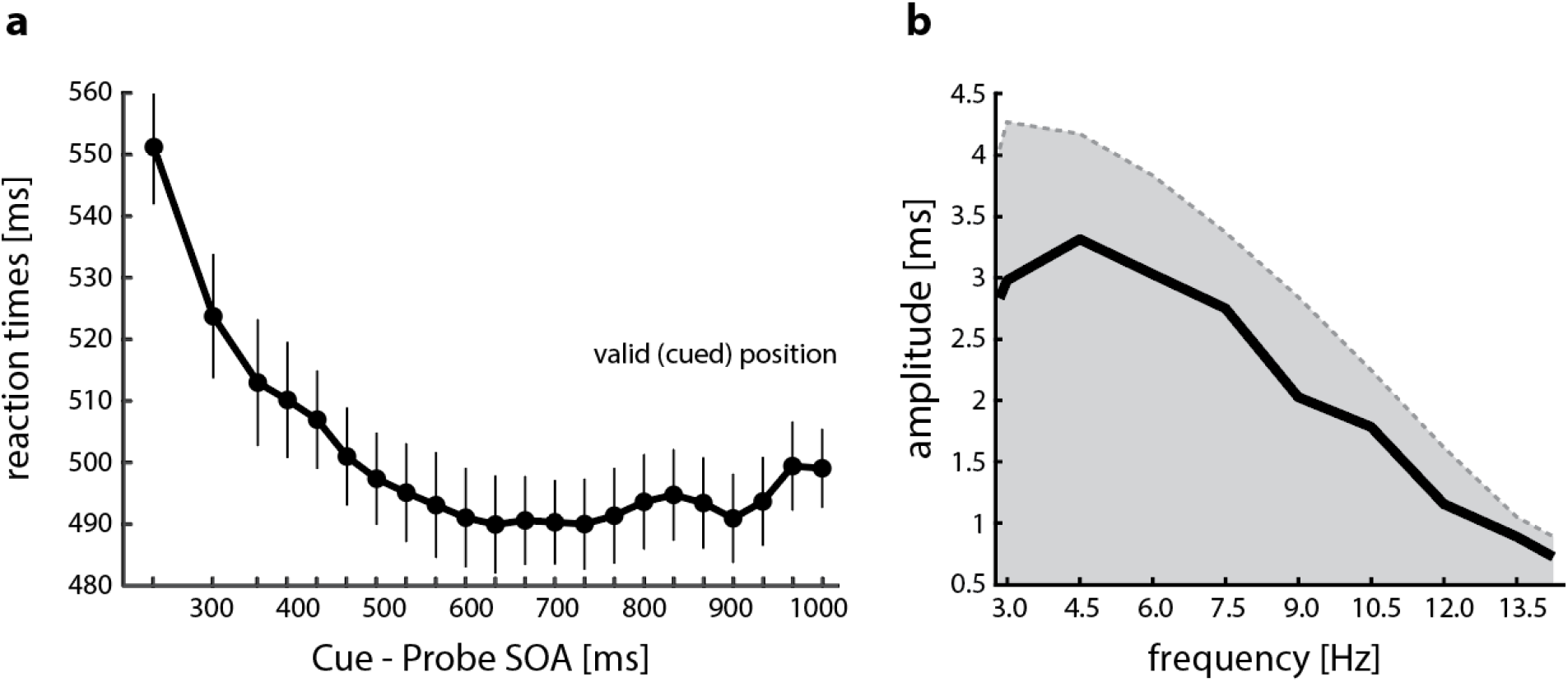
Temporal dynamics of RTs to probes at the cued memory position. **a** Reaction times (of correct responses) to the probe at the cued memory position (valid trials) as a function of time after the onset of the attentional cue (SOA: stimulus-onset-asynchrony). Vertical lines indicate standard error of the mean. **b** Fourier amplitude spectrum of the time-course of RTs to probes at the cued position. Shaded area represents the 95^th^ percentile of the permutation null-distribution.

**Supplementary Figure 5.**
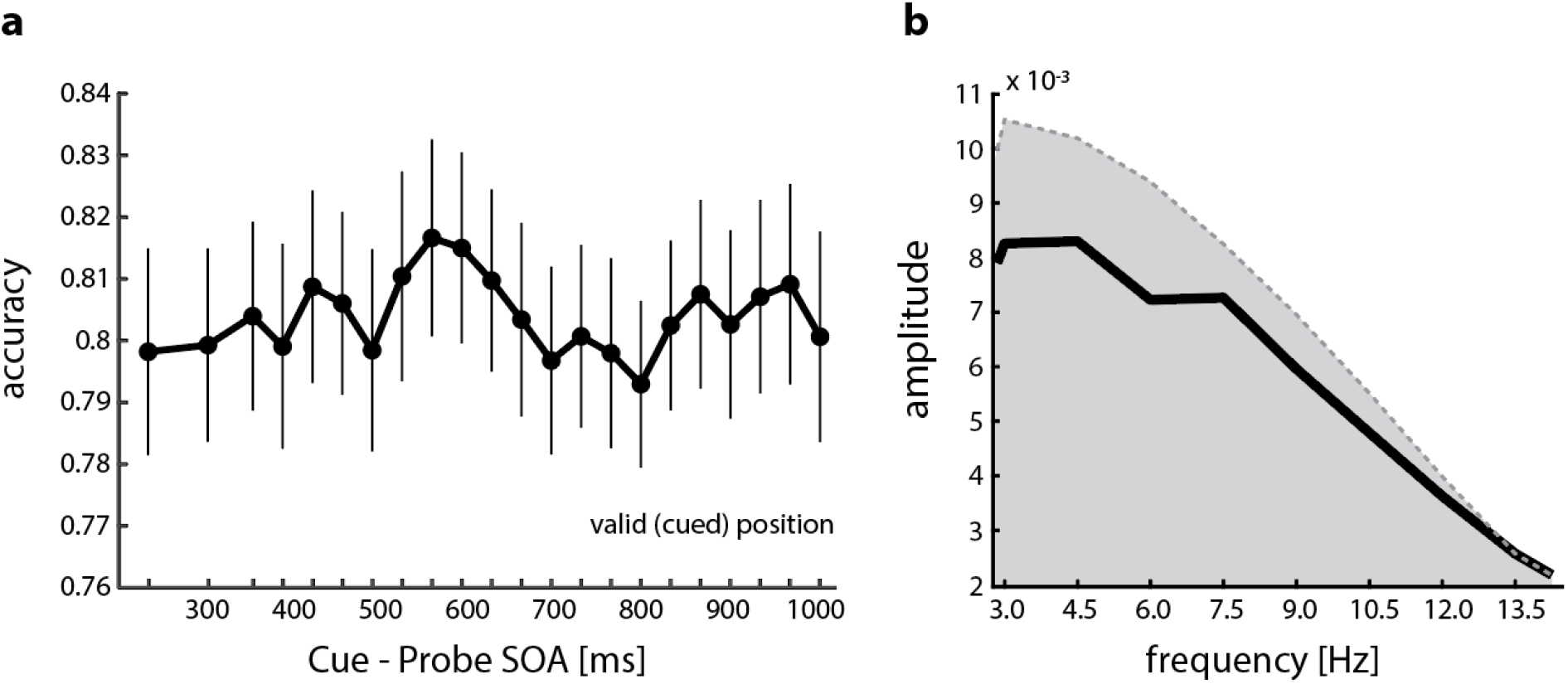
Temporal dynamics of accuracy to probes at the cued memory position. **a** Accuracy of responses to the probe at the cued memory position (valid trials) as a function of time after the onset of the attentional cue (SOA: stimulus-onset-asynchrony). Vertical lines indicate standard error of the mean. **b** Fourier amplitude spectrum of the time-course of accuracy of responses to probes at the cued position. Shaded area represents the 95^th^ percentile of the permutation null-distribution.

## Footnotes

1 In the perceptual literature, oscillations of performance are often discussed in the context of ‘rhythmic attentional sampling’. The term sampling may imply that attention periodically fully switched between objects. We therefore prefer to use the term “attentional prioritization” in the context of our results.

